# *In vitro* sexual dimorphism establishment in schistosomes

**DOI:** 10.64898/2026.01.20.700612

**Authors:** Rémi Pichon, Magda E Lotkowska, Jude L. D. Bulathsinghalage, Madeleine McMath, Mary Evans, Benjamin J. Hulme, Kirsty Ambridge, Geetha Sankaranarayanan, Simon Kershenbaum, Sarah D. Davey, Josephine E. Forde-Thomas, Karl F. Hoffmann, Matthew Berriman, Gabriel Rinaldi

**Author notes:** Correspondence: co-corresponding authors: Matt Berriman, Gabriel Rinaldi. co-first authors (Rémi Pichon, Magda E Lotkowska).

## Abstract

Schistosomes are parasitic flatworms that cause Schistosomiasis, a major Neglected Tropical Disease (NTD) that affects more than 250 million people worldwide. With two morphologically distinct sexes, a heterogametic female (ZW) and a homogametic male (ZZ), schistosomes are an exception among flatworms, which are largely hermaphroditic. Sexual dimorphism in schistosomes only becomes apparent by adulthood within the mammalian host. The cellular and molecular mechanisms underlying the sexual differentiation of these parasites are poorly understood, partly due to intrinsic challenges in assessing parasite development *in vivo*. Therefore, robust and reproducible approaches for maintaining and developing parasites *in vitro* are needed to overcome these difficulties. However, to date, only a few studies have focused on protocols that allow cultured parasites to reach sexual dimorphic stages, and none have been reproduced, limiting the ability to understand the unique sexual biology of this major human parasite. Here, we refine a protocol for long-term culture of newly transformed cercariae that developed *in vitro* into sexually dimorphic forms. We assessed the effect of two different sera, Foetal Bovine Serum (FBS) and Human Serum (HS), added to the culture medium supplemented with human red blood cells. We found that in contrast to FBS-cultured worms, HS-cultured parasites digested red blood cells, a crucial step for long term parasite development. Additionally, while most FBS-cultured parasites did not progress beyond an early liver stage, sexual dimorphism was clearly established in the HS-cultured worms, albeit delayed compared to *in vivo* development. Moreover, EdU pulse experiments revealed a continuous proliferation of cells over time in HS-cultured parasites, while a significantly lower number of proliferating cells were detected in FBS-cultured worms. This protocol paves the way to study parasite development *in vitro*, positively impacting the principles of the 3Rs (Replacement, Reduction and Refinement) for animal research, and allowing for in-depth studies of sexual dimorphism establishment as well as *in vitro* screening for novel control strategies across the life cycle of these major human parasites.

## Introduction

Schistosomiasis, a major Neglected Tropical Disease (NTD) affecting >250 million people worldwide, is caused by the infection with blood flukes (class Trematoda) in the genus *Schistosoma*^1^. The pathology associated with schistosomiasis is largely driven by egg trapping in different tissues depending on the species. Infection with *Schistosoma mansoni* leads to eggs lodged mainly in the liver, inducing inflation, fibrosis and granuloma formation^2^. This suggests that interfering with the sexual development of schistosome intra-mammalian stages could potentially restrict human pathology. Currently, the complete reliance on a single drug (Praziquantel) to treat the infection and its use in drug mass administration programs in endemic areas threatens the development of drug resistance^3,4^. Therefore, novel control strategies are urgently needed, and identifying original targets for drug/ vaccine development became a priority. A better understanding of the mechanisms underlying schistosome development, including sexual dimorphism establishment, will pave the wave to achieve this goal.

While most parasitic flatworms are hermaphrodites, schistosomes are dioecious with genetically determined female (2n=16, ZW) and male (2n=16, ZZ) individuals. Whilst male and female parasites are morphologically indistinguishable across most developmental stages, sexual dimorphism becomes apparent by adulthood within the mammalian host^5^. Male and female worms undergo separate but concurrent sexual differentiation of their gonads and somatic tissues that eventually allows intersexual pairing, a critical step for female maturation, egg production and life cycle propagation^6,7^. Transcriptomic studies, at both bulk^8–12^ and single cell^13–15^ levels for intra mammalian stages *in vivo* and *ex vivo*, have revealed key pathways and molecular mediators underlying the critical process of female sexual maturation induced by pairing. However, the cellular and molecular mechanisms driving sexual dimorphism prior to pairing remain poorly characterised, partly due to intrinsic challenges in assessing the development of the parasite *in vivo*^12^.

Approaches to better understand the biology of parasites at large have been developed and optimised using *in vitro* or *ex vivo* refined culture systems^16,17^. Combinations of culture media components, sera from different sources and additives have been tested^18,19^. In addition, *organ-on-a-chip* systems and organoid-based 2D and 3D culture platforms^20,21^ have first been implemented for protozoa parasites such as *Plasmodium*, *Toxoplasma*, and *Cryptosporidium* species. These methods have facilitated the molecular dissection of processes underlying single-cell parasite development and interaction with the host^22–24^. Recently, these technologies have started to be transferred to metazoan parasites to study host-parasite interaction and immune response in the context of helminth infections^25–29^. *In vitro* cultivation of both larval tapeworms (cestodes) and stem cells isolated from cestodes, coupled with ‘omic’ and imaging analyses have unveiled the role of stem cells in parasite development^30,31^. However, despite these significant developments in the field, reproducing the natural development of trematodes under controlled *in vitro* conditions remains extremely challenging^17^.

Significant progress was achieved in culturing schistosomes when, more than forty years ago, Paul F. Basch developed and optimised comprehensive culture protocols for intra-mammalian developmental stages of schistosomes^32–34^. Basch reported for the first time the development of male and female parasites from cercariae, with worm pairing and production of infertile eggs entirely *in vitro*^32^. Remarkably, to the best of our knowledge, these results have not been replicated and no further attempt to obtain egg-laying worm pairs developed *in vitro* from cercariae has been reported since Basch’s seminal studies^17^. More recently, *in vitro* culture protocols for schistosomes have been refined either to maintain *ex vivo* parasites collected from infected mice to study parasite reproduction biology^35,36^, or to culture juvenile parasites from cercariae and develop drug screening approaches^37–39^. However, no long-term culture conditions to specifically assess schistosome sexual differentiation have been reported since 1981^33^. Despite one research group referring to the approach^40^, there has been no detailed description and no widespread adoption. In fact, published culturing studies have continued to utilise FBS^35,41^.

Novel functional approaches applied to culture systems that allow reliable and reproducible *in vitro* establishment of sexual dimorphism will shine new light into molecular mechanisms underlying schistosome intra-mammalian development. We aimed at optimising a platform to study intra-mammalian schistosomes that supports *in vitro* sexual dimorphism establishment and consequently leading to an overall positive impact in the 3Rs (Reduction, Replacement, Refinement) on animal research (https://nc3rs.org.uk/)^42^. Here, we refined a protocol for long-term culture of newly transformed cercariae by assessing the effects of two different sera, Foetal Bovine Serum (FBS) and Human Serum (HS), added to the medium supplemented with human Red Blood Cells (hRBCs). Striking differences were evident between parasites maintained in either FBS or HS. First, phenotypic differences between FBS- and HS- cultured parasites became evident as early as 48 hours in culture, with HS-cultured parasites exhibiting higher rates of cell proliferation resulting in larger worms in the HS condition. Second, hRBC were digested and hemozoin became apparent in the intestines of HS-cultured parasites within a day in contrast to FBS-cultured parasites that mostly lacked visible hemozoin. Finally, while most of the FBS-cultured parasites did not progress beyond lung and early liver stage, HS-cultured parasites reached sexually dimorphic stages by week 6, albeit at a slightly delayed rate compared to *in vivo* development. In the mouse model, parasites become dimorphic by day 21 post-infection (∼3 weeks)^12^. Taken together, this protocol allows early sexual development of schistosomes to be studied *in vitro*, provides a method for high throughput drug screening targeting parasite development, and reduces the reliance on mammalian hosts in research.

## Results

### Sexually dimorphic schistosomes developed entirely *in vitro* from cercariae

Searching for culture media able to cultivate parasites from cercariae to maturity, we decided to test Human Serum (HS) compared to the commonly used Foetal Bovine Serum (FBS). This rationale was based both on the fact that humans are the definitive host of *Schistosoma mansoni*, and that two previous reports in 1981 showed that HS was capable of producing mature schistosomes *in vitro*^32,33^, but they were never fully replicated. More recently, culture media were complemented with HS in studies focused on early stages of parasite development for drug testing^37–39^; however, these worms never progressed beyond the liver stage^43^. Therefore, to ascertain the *in vitro* sexual dimorphism establishment of schistosomes entirely developed from cercariae, we compared the effect of two different sources of blood serum: FBS and HS. The development of schistosomula derived from mechanically transformed cercariae was assessed in at least 15 independent experiments, five of which were maintained over a period of at least 10 weeks to assess parasite survival and ability to mate and produce fertile eggs (**Figure 1A; Supplementary Table S1**).

**Figure 1.**
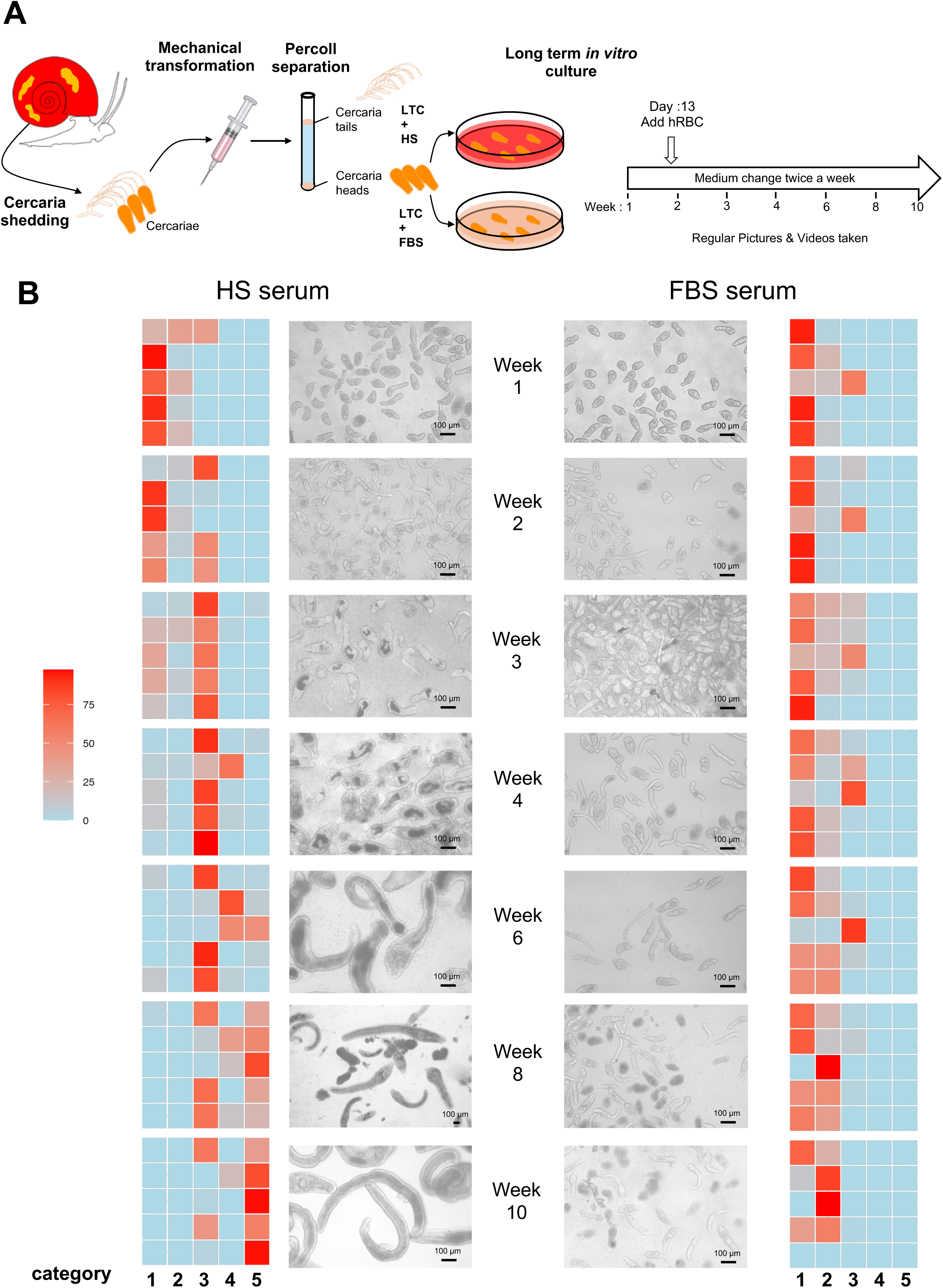
Dimorphic female and male schistosomes entirely developed in vitro from cercariae. **A.** Schematic representation of the collection and mechanical transformation of cercariae into schistosomula for long-term *in vitro* culture. **B.** Morphological scoring of cultured *S. mansoni* schistosomula at the indicated time points after *in vitro* transformation (week 1 to week 10) for worms in Long Term Culture (LTC) medium supplemented either with Human Serum (HS - left) or Foetal Bovine Serum (FBS - right). Heatmap columns represent five distinct morphological categories, and rows indicate 5 independent culture experiments, i.e. parasites obtained from different batches of infected snails. Heatmap colours represent the percentage of worms for each replicate in each morphological category. Middle panel: Representative images of *in vitro* schistosomula cultured in either HS or FBS as indicated. Scale bars: 100 µm. Category 1: Early schistosomula; Category 2: Lung schistosomula; Category 3: Early liver schistosomula; Category 4: Late liver schistosomula; Category 5: Dimorphic schistosomula. A detailed description of the developmental categories and representative images are provided in Supplementary Figure S1.

No morphological differences were observed between parasites cultured either in FBS or HS within the first week in culture; in both conditions most parasites were classified as early schistosomula [category 1: 76% ± 30 (average ± SD) in FBS and 73% ± 29 (average ± SD) in HS] with few lung (category 2) and early liver schistosomula (category 3) (**Figure 1B, week 1; Supplementary Figure S1**). The mean mortality (category 0) at week 1 was slightly higher, but not statistically significant (P= 0.42), in worms cultured in HS [9.75% ± 2.76 (average ± SD)] compared to the mortality registered in FBS-cultured parasites [5.52% ± 5.18 (average ± SD), **Supplementary Table S6**], consistent with previous findings^39^.

Differences in parasite development between the two conditions became apparent by week 2 (**Figure 1B**). At this time point, 14.8% ± 24.9 (average ± SD, excluding dead worms) or 36% ± 33.6 (average ± SD, excluding dead worms) of the parasites cultured in FBS or HS, respectively, have reached category 3, i.e., early liver schistosomulum. Parasites in FBS rarely progressed beyond this stage during the 10-week experiment, with very few parasites (<0.1% ± 0.2, average ± SD) reaching category 4, i.e., late liver schistosomulum. In contrast, worms cultured in HS developed over time across all categories, achieving marked sexual dimorphism by week 6 (13.4% ± 18.6, average ± SD) (**Figure 1B; Supplementary Figure S3A**), as confirmed by PCR (**Supplementary Figure S3B; Supplementary Table S2**). No differences in the timing for sexual dimorphism establishment were observed between male and female parasites. The mortality rate of FBS-cultured parasites reached an average of 76.24% ± 23.46 (average ± SD) by week 10, after which the experiments under this condition were stopped as most parasites were dead (**Supplementary Figure S2**). From that time point onwards only parasites in HS were kept in culture. As previously described for the *in vivo* development of schistosomes^12^, *in vitro* cultured parasites showed developmental asynchrony in agreement with Basch’s observations^33^; however, by week 10 most of the worms in HS (73.7% ± 25.4, average ± SD) acquired an evident sexual dimorphism (**Figure 1B**).

The differences in parasite development between the two tested conditions were also confirmed by measuring worm areas at different time points in culture (**Figure 2; Supplementary Table S3**). Parasites cultured in the presence of HS grew exponentially over time to reach on average 110,069 μm^2^ ± 72,959 (average ± SD), ∼20-fold larger than parasites cultured in FBS for 10 weeks, which only grew slightly, plateauing at an average 5,417 μm^2^ ± 1,322 (average ± SD), similar to the size reached by parasites after only two weeks in culture with HS (P-value < 0.01, **Supplementary Table S6; Figures 1 and 2**).

**Figure 2.**
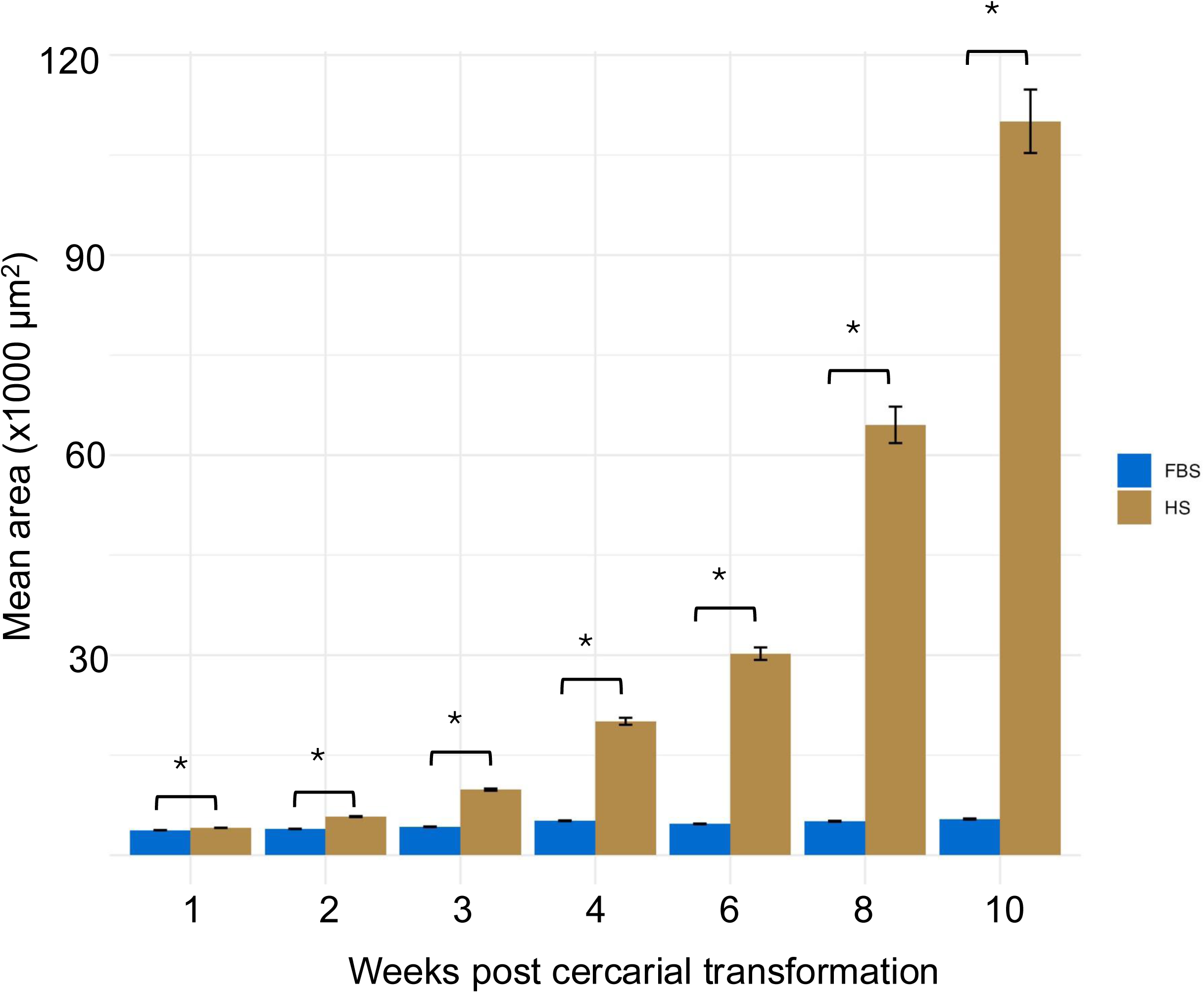
Parasites cultured in Human Serum grew in size unlike those cultured in Foetal Bovine Serum. Bar plot representing area measurements of schistosomula developed *in vitro* and cultured in media complemented either in Foetal Bovine Serum (FBS) or Human Serum (HS) at indicated weeks after cercarial transformation. Bars indicate mean area (µm²) ± SEM. HS: Human Serum (light brown); FBS: Foetal Bovine Serum (blue). *: P-value < 0.01 in indicated pairwise comparisons (Supplementary Table S6).

### Parasites developed in human serum readily digest red blood cells

It is widely accepted that within the mammalian host, schistosomes begin to feed on blood cells through their digestive tract at 10 days post-infection; at this time the majority of the parasites have left the lungs and started to colonise the portal system veins in the liver^44^. Based on both previous reports^45^, and pilot experiments in which adding human Red Blood Cells (hRBCs) to the culture before day ∼10 did not show obvious haemoglobin digestion, we decided to supplement the culture media with hRBCs at day 13. The addition of hRBCs allowed the parasites to feed and thus continue their development^19^. At this point, they began to swallow and degrade erythrocytes, producing hemozoin, a black pigment derived from host haemoglobin degradation and visible in the worms’ intestines. Even though few parasites in FBS reached the early liver stage (category 3) within the first week in culture, a minority of them were able to digest hRBCs (3.6% ± 4.7, average ± SD), indicated by displaying black guts (BG) (**Figure 3**). In contrast, a significant proportion of HS-cultured parasites (36.2% ± 33.6, average ± SD) had already reached the early liver stage by the second week in 3 out of 5 experiments (**Figure 1B**, week 2). Moreover, parasites cultured in HS displayed a functional digestive system capable of assimilating hRBCs; more than half of the HS-cultured parasites (65% ± 6, average ± SD, P-value < 0.05) showed BG in comparison to FBS-cultured parasites (3.7 ± 4.7 %) (**Figure 3; Supplementary Table S4**).

**Figure 3.**
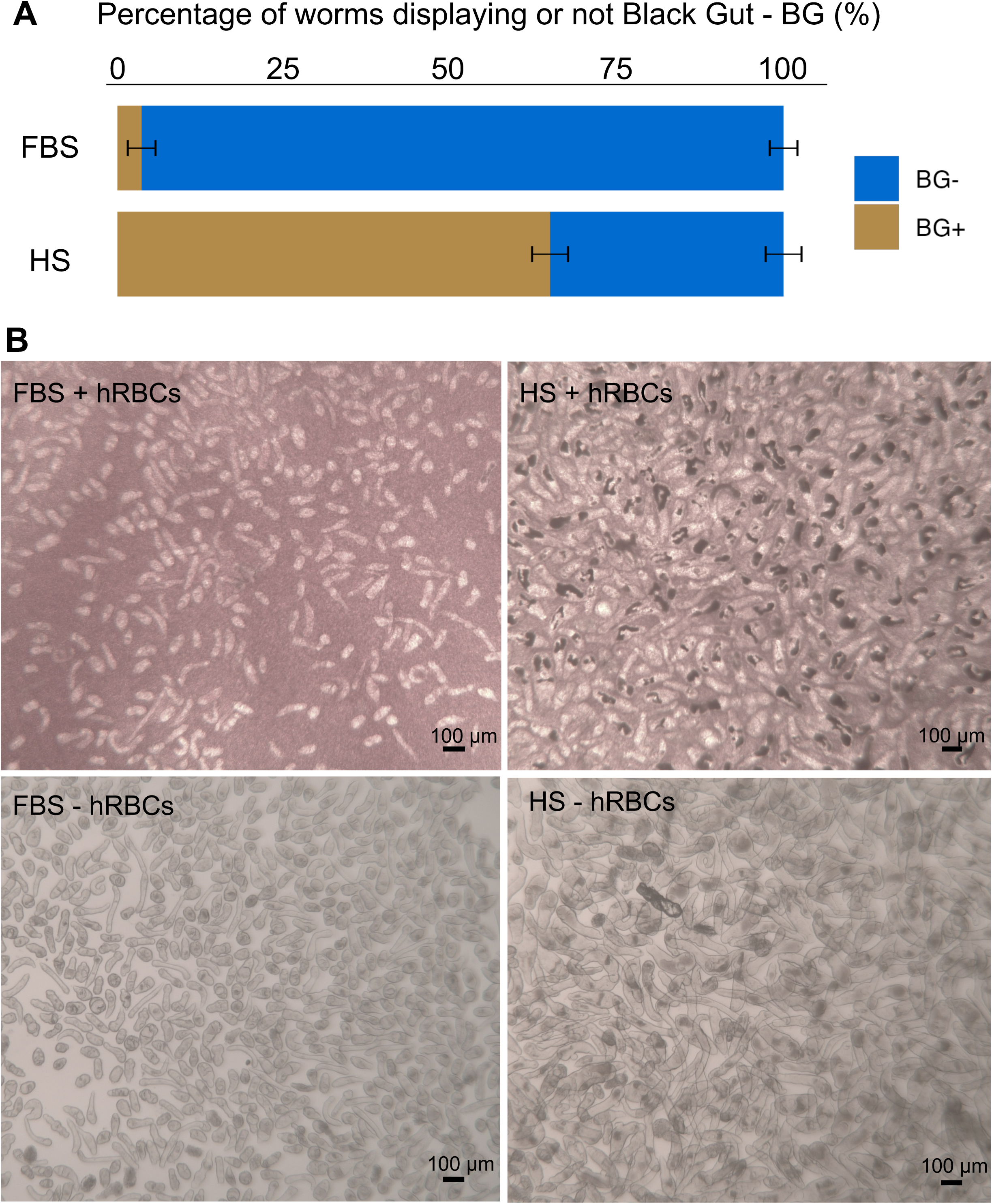
Parasites developed in Human Serum readily digest red blood cells. **A.** Bar Plot representing the percentage of Human Serum (HS)- or Foetal Bovine Serum (FBS)-cultured schistosomula with (blue bar) or without (light brown bar) black guts (BG) due to the presence of intestinal hemozoin. Washed human Red Blood Cells (hRBCs) were added into the media at day 13 post-transformation and images captured one or two days later. Error Bars = SEM. (Statistical analyses in Supplementary Table S6.) **B.** Representative images of in vitro developed schistosomula cultured in FBS or HS one day after adding hRBC (+ RBC) and controls without RBC (- RBC). Scale bars: 100 µm.

### Development of parasites in human serum may be driven by stem cell proliferation

The growth and development of an organism is driven by a finely regulated combination of cell hyperplasia, hypertrophy, and apoptosis across different tissues^46^. In schistosomes, a complex stem cell system consisting of both somatic and germline stem cells has been described by leveraging recent single cell transcriptomic data across different developmental stages, including schistosomula and adult worms^47^. Therefore, we decided to investigate whether the differences in development, growth and feeding capacity between parasites cultured in either FBS- or HS- complemented medium were associated with distinct cellular proliferation rates. EdU pulse experiments revealed notably lower cell proliferation in FBS-cultured parasites as early as 2 days post transformation compared to HS-cultured parasites (**Figure 4**). It has previously been demonstrated that a group of 5 well-defined somatic stem cells is the only set of cells that actively proliferate in 2-day schistosomula^48^. Hence, the proliferating cells observed and quantified in our study were most likely stem cells (**Supplementary Figure S4; Supplementary Video S1**). The difference between the number of proliferating stem cells in parasites cultured in FBS or HS increased significantly over time. HS-cultured schistosomula showed higher numbers of proliferating stem cells, with a median of >48 and >60 EdU+ cells per worm at days 8 and 15, respectively (**Figure 4**). On the other hand, most FBS-cultured parasites displayed no more than an average of 20 EdU+ cells per worm (**Figure 4**). hRBCs were added at day 13 post-cercarial transformation to both FBS- and HS- complemented culture media. Worms kept in FBS or HS in the absence of hRBCs were included as controls. Regardless of the serum employed, no significant differences in the numbers of proliferating cells were observed between worms cultured in the presence or absence of hRBCs (**Supplementary Tables S5 and S6**).

**Figure 4.**
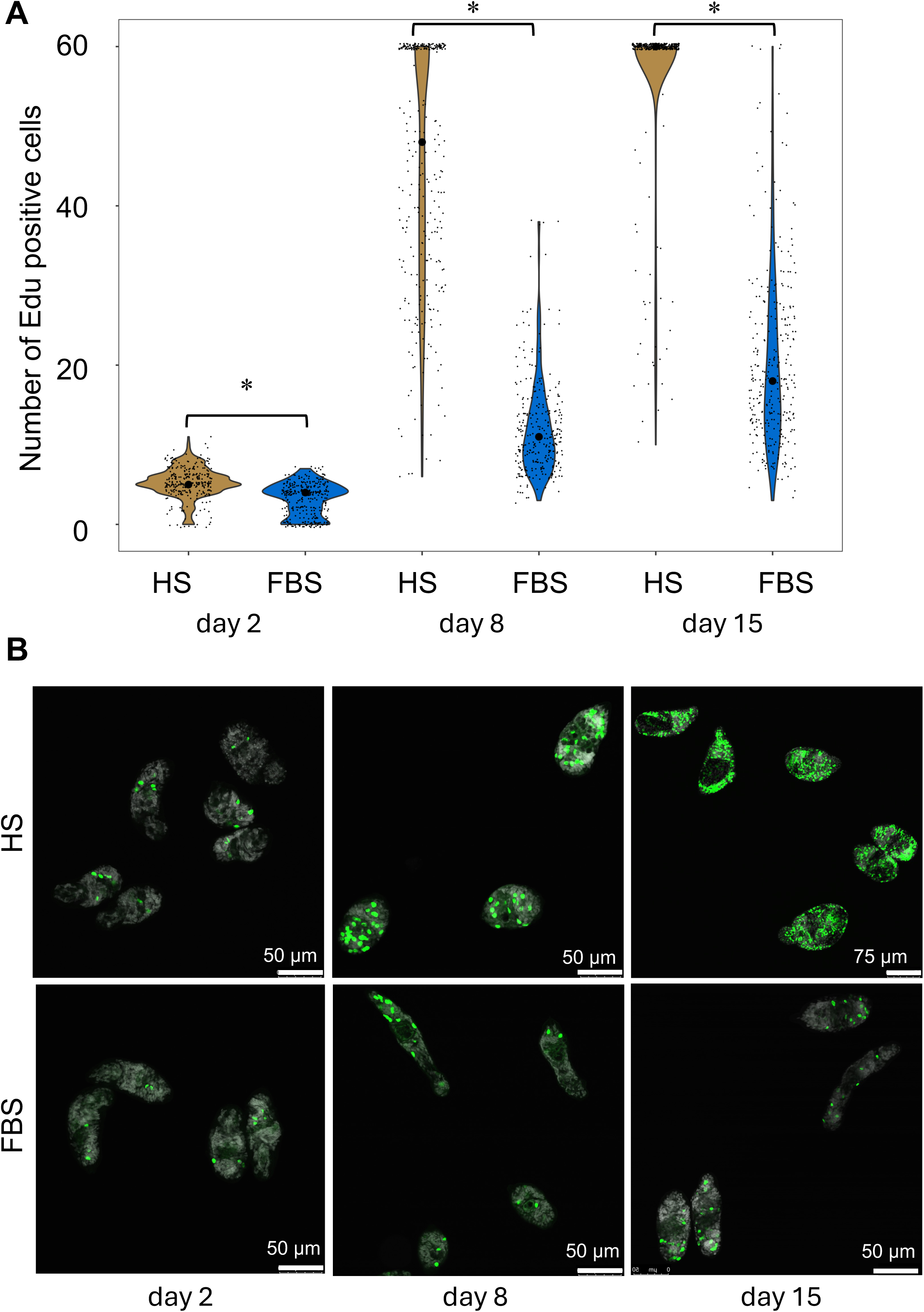
Development of parasites in human serum may be driven by stem cell proliferation. **A.** Violin plots showing the number of EdU+ cells per worm at indicated time points (2, 8, and 15 days post cercarial transformation) in parasites cultured either in Foetal Bovine Serum (FBS, blue) or Human Serum (HS, light brown). Human Red Blood Cells (hRBCs) were added in the culture at day 13 post cercarial transformation. The small black dots indicate individual worms, and the big black point indicates the median of EdU+ cells per worm. All worms showing Ill 60 EdU+ cells were counted and clustered together in the group named ‘60 EdU+ cells’. Hence, the data were treated as ordinal, and statistical analysis performed by Kruskal-Wallis test with Dunn multiple comparison post-hoc test, with P≤0.05 (*) considered significant (Supplementary Tables S5 and S6). **B.** Representative images of parasites displaying Edu+ cells at each indicated time point and culture condition. Edu+ cells and nuclei were labelled with Alexa fluor 488 (green) and DAPI (white/grey), respectively. Scale bars: 50 µm or 75 µm as indicated.

### *In vitro* cultured schistosomes display sexual dimorphism, developing reproductive systems and pairing capacity

*In vitro*-developed schistosomes began to show sexually dimorphic features from day ∼42 onwards in HS-supplemented culture medium (**Figure 1**, week 6). Confocal microscopy of DAPI- and Phalloidin- stained *in vitro* cultured male worms confirmed the presence of the gynaecophoric canal, developing 3 to 5 testis lobes with sperm cells, and cirrus (**Figure 5A-D; Supplementary Video S2**). Likewise, we confirmed the presence of primordial ovaries, oviduct, ootype and uterus in female parasites entirely developed *in vitro* (**Figure 5E-H; Supplementary Video S3**). In some experiments sexually dimorphic parasites cultured in HS medium were kept alive for more than 150 days (**Supplementary Figure S5**). While the establishment of sexual dimorphism was robust and reproducible in more than 15 independent experiments, pairing between male and female parasites was rare. Pairing was observed only in experiments lasting more than 80 days in which we were only able to observe a few couples pairing up, and in addition, this pairing was temporary (**Figures 6A, 6B; Supplementary Video S4**).

**Figure 5.**
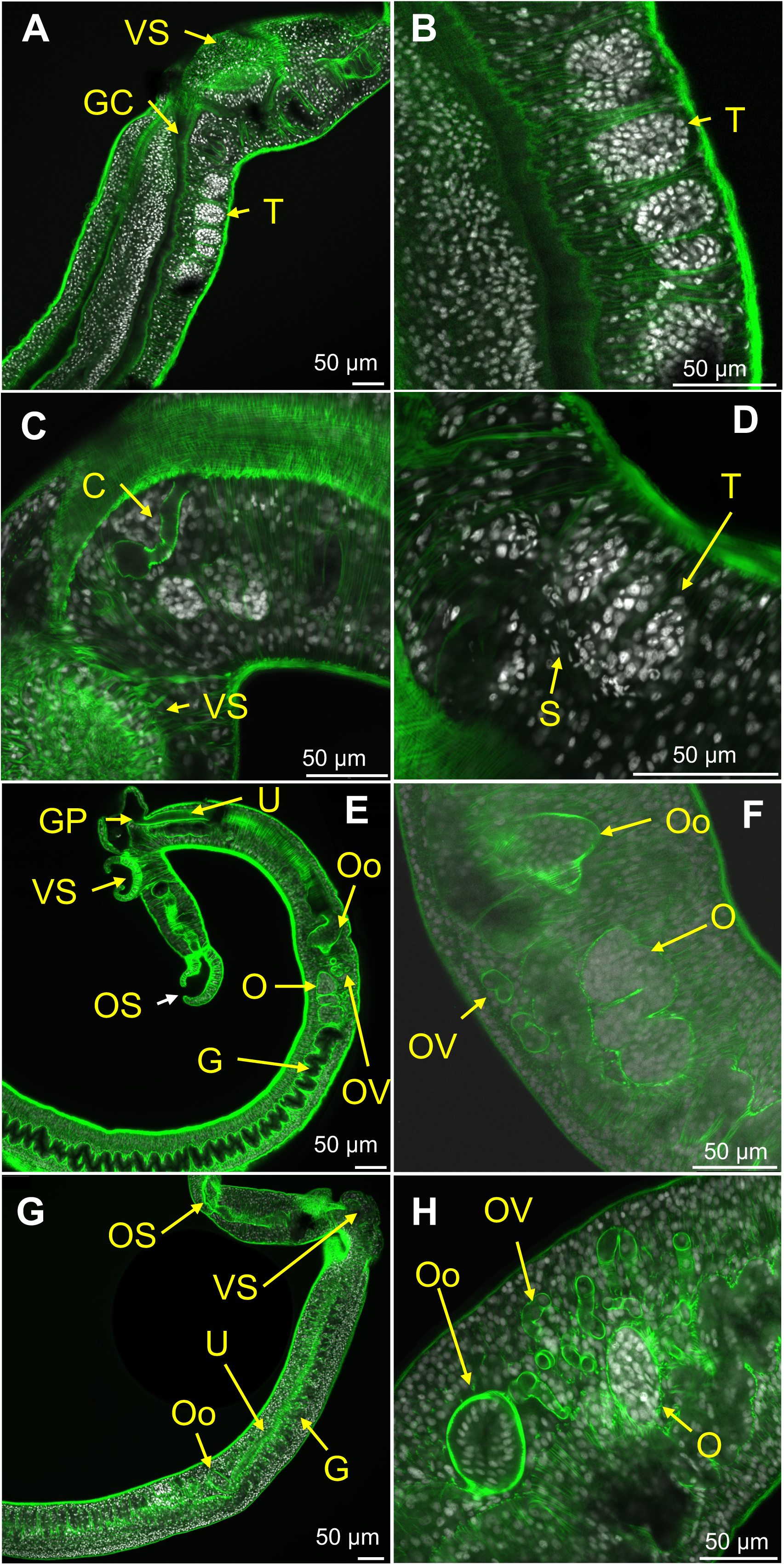
*In vitro* cultured schistosomes display sexual dimorphism and developing reproductive systems. Representative confocal microscopy images of *in vitro* developed male (A-D) and female (E-H) at day 60 in culture. Worms shown in panels G and H are different individuals. CellMask Green Actin Tracking Stain: green, DAPI: grey (A-E, G), cyan (F, H). C: Cirrus. T: Testis, OS: Oral sucker. VS: Ventral sucker. GC: Gynaecophoric canal. G: Gut. GP: Genital pore. U: Uterus. O: Ovary. Oo: Ootype. OV: Oviduc. S: sperm. Scale bar: 50 µm.

**Figure 6.**
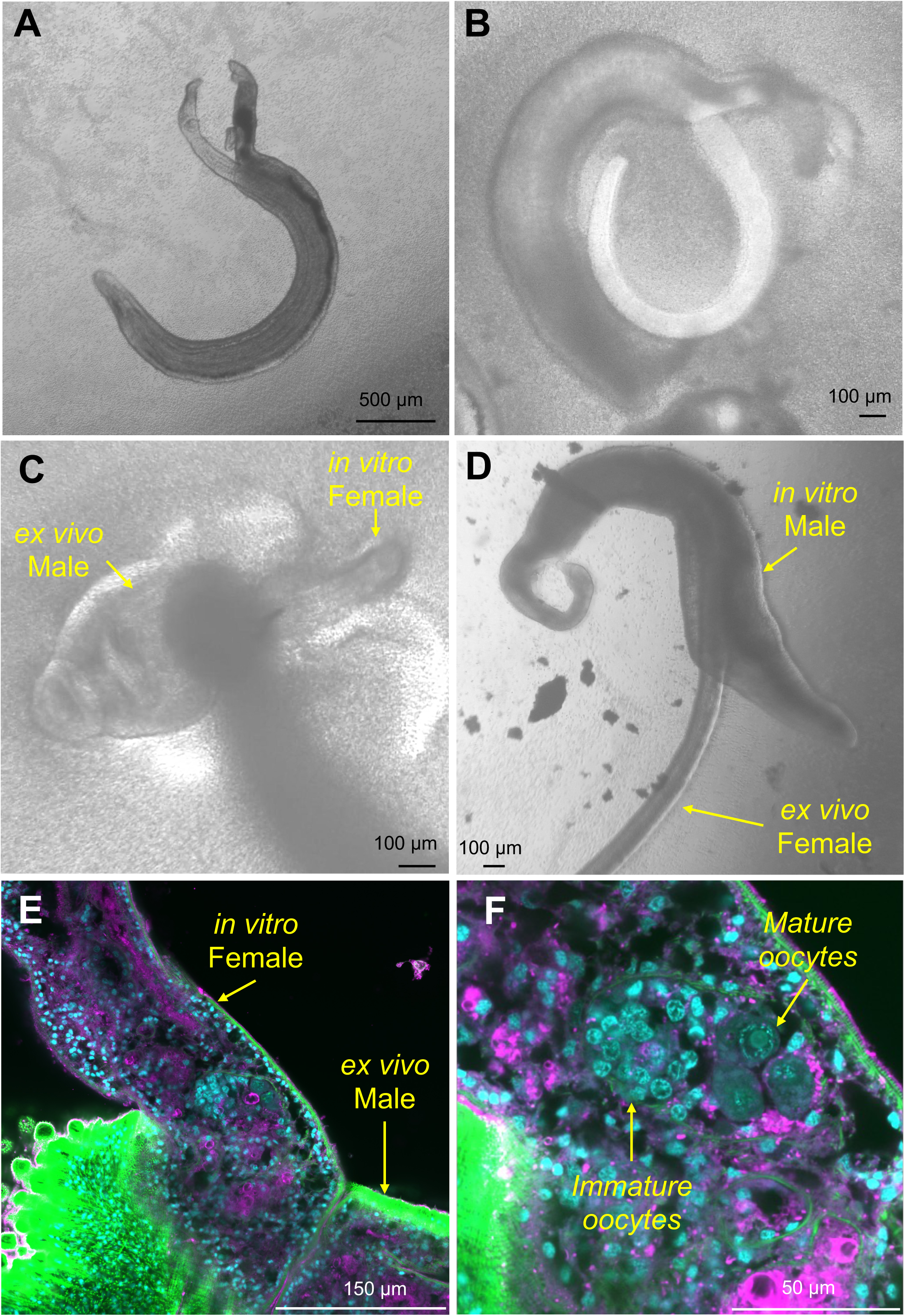
*In vitro* cultured schistosomes are capable of pairing. **A, B.** Representative bright-field images of pairs of schistosomes developed entirely *in vitro* after 80 (A) and 150 (B) days of culture in culture medium supplemented with HS. Scale bars: 500 µm (A), 100 µm (B). **C, D.** Representative bright-field images of a worm pairs between *ex vivo*-collected males and *in vitro*-developed females (C) and *ex vivo*-collected females and in vitro-developed males (D) within 24 hours after placing the worms in the same well to facilitate pairing. Scale bars: 100 µm, **E, F.** Confocal microscopy image of a schistosome pair (*ex vivo*-collected male and *in vitro*-developed female) in copula (E), and magnification of the ovarian area, highlighting maturing oocytes (F). CellMask Green Actin Tracking Stain: green, DAPI: grey, CellMask Deep Red Plasma Membrane Stain: magenta. Scale bar: 150 µm (E), 50 µm (F).

Considering the rarity of the pairing between *in vitro* developed male and female worms, we investigated whether these parasites display the capacity of pairing with *in vivo* developed worms collected from experimentally infected mice. Male and female adult worms were recovered from mice by portal perfusion on day 42 post-infection, sorted by sex and placed in culture with worms of the opposite sex developed *in vitro*. Within 24 hours of initiating the co-culturing of *in vitro* developed worms with *ex vivo* collected worms, couples were observed (**Figure 6C, D; Supplementary Video S5**). These findings suggested that the *in vitro* developed and sexually dimorphic parasites were capable of intersexual pairing. Moreover, *in vitro* developed females coupled with *ex vivo* collected mature males displayed signs of primordial ovary maturation with larger oocytes towards the posterior region of the ovary (**Figure 6E, F; Supplementary Video S6**). On the other hand, females developed *in vitro* but not paired with *ex vivo* males remained immature. Remarkably, in more than 30 independent *in vitro* culture experiments, where male and female parasites developed sexual dimorphism and eventually paired up, no eggs were produced or laid. This indicates that further refinements in the culture protocol are needed to advance parasite development (the *in vitro* development was delayed compared to the *in vivo* development), increase the likelihood of pairing, and facilitate the production of eggs.

## Discussion

Recent progress in culture systems and functional genomic tools is allowing researchers to address long standing questions in helminth biology that were previously inaccessible to experimentation^49–53^. Among these enduring questions, stand those related to the unique sexual biology of the trematode family Schistosomatidae. Schistosomes are dioecious with genetically determined female and male individuals, which is unusual for flatworms^5,54^. However, the sexual dimorphism of male and female schistosome worms only becomes established within the mammalian host, and represents a critical step towards worm maturation, intersexual pairing, and egg production^55^. A better understanding of the molecular and cellular basis underlying sexual dimorphism establishment in schistosomes would lead to approaches to block parasite development and life cycle propagation. Studying the sexual development of schistosomes by performing controlled experiments in which parasites can be co-cultured, genetically manipulated or treated with different compounds requires robust and reproducible protocols for long-term *in vitro* culture. Culture protocols have recently been developed to maintain *ex vivo* schistosomes collected from infected mice^35,36^, or to obtain juvenile parasites from cercariae for drug screening experiments^37–39,56^. However, no long-term culture conditions have been successfully refined to experimentally assess sexual differentiation and dimorphism establishment in schistosomes. Here, aiming to refine a culture medium formulation that supports *in vitro* parasite development and establishment of sexual dimorphism, we compared the effect of modified Basch’s media^19^, supplemented with human Red Blood Cells (hRBCs) and 20% of either Fetal Bovine Serum (FBS) or Human Serum (HS). While initial parasite survival and development appeared comparable in both conditions during the first week in culture, striking morphological differences emerged from week 2 onwards. Parasites cultured in HS, progressed through all developmental categories, and acquired sexual dimorphism by week 6. On the other hand, parasites maintained in FBS were stunted at early stages (mainly lung stage). These experimental outcomes were consistent with the findings reported by Paul F. Basch^32^ in 1980. Probably, from the 1980s onwards a combination of factors that include ethical concerns, variability associated with human-derived products and safety considerations related to blood borne viruses such as HIV, and Hepatitis B and C, determined that parasitologists favoured the use of FBS over HS for helminth culture. FBS has been extensively used in cell culture media, refined and adapted to diverse human and animal cell types since 1958^57^ providing a reliable source of amino acids, carbohydrates, hormones, lipids, proteins, vitamins, and growth factors^58^. Nevertheless, the scientific community is increasingly advocating for the replacement of FBS as a supplement in tissue culture^59^. This trend is driven by limitations associated with FBS, including the presence of undefined tentatively harmful factors for the cells in culture, lack of reproducibility, lack of transparency in its production, and critically ethical concerns related to animal welfare^60^. Our protocol, that supports the fully *in vitro* development of schistosomes, would also positively impact the 3Rs (i.e., Reduction, Replacement, Refinement)^42^ for minimising the use of animals for research and serum production^61,62^.

Parasites cultured in the presence of HS not only developed into sexually dimorphic male and female worms, but strikingly, were able to digest hRBCs and process haemoglobin within one day after the addition of the cells. These findings may reflect differences in gastrodermis and oesophageal gland development; worms cultured in the presence of HS displayed a differentiated gastrodermis that enabled haemoglobin digestion and accumulation of hemozoin within the intestines^63^. In contrast, most parasites cultured in FBS were unable to produce hemozoin in the presence of hRBCs. This may be due to impaired gastrodermis differentiation that ultimately would lead to stunted development and death. Hematophagous parasites including Plasmodium species and schistosomes, obtain key nutrients via the proteolysis of host hemoglobin. However, this process leads to the production of free-heme groups which in turn generate highly toxic oxygen free radicals and lipid peroxidation^64^. These toxic free-heme derivatives become inactive when aggregated into an inert crystalline polymer, named hemozoin, observed as dark pigment within the intestines of schistosome intra-mammalian stages^63^. In addition, increasing evidence shows that hemozoin may play critical roles during parasite development by supplying iron for egg production^63^, and interaction with the mammalian host by an immune modulatory role^65^. In the mouse model, schistosomula begin to feed on blood once they have left the lungs and reached the portal system 9 to 11 days post infection^44,66^. Moreover, schistosomula collected from experimentally infected mice at day 13 post infection already show hemozoin in their guts^12^. Similarly, HS-developed schistosomula were able to feed on blood *in vitro* within a comparable time window; hRBCs were added in the medium on day 13 in culture, and within 24 hours most of the parasites’ gut contained hemozoin. These findings suggest that our *in vitro* culture system successfully recapitulates *in vivo* gastrodermis development of schistosomula. A better understanding of the *in vitro* development of the parasite oesophageal gland and gastrodermis and its role in the production of hemozoin will expose tentative novel targets for control^67^.

Although previous reports have shown that it is possible to cultivate parasites in cell-free environments^39,43^, adding hRBC has proven essential for long-term maintenance of parasites and for establishing sexual dimorphism. EdU pulse experiments suggested that schistosome *in vitro* development in the presence of HS may be driven by the proliferation and differentiation of stem cells. The stem cell system in schistosomes has been extensively studied since the first description of somatic stem cells in adult worms^68^, followed by single cell transcriptomic identification and functional characterisation of three key stem cell populations in intra-snail stages^48^. Two of these stem cell types, maintained in intra-mammalian stages, proliferate and differentiate into precursors of somatic and germ line cells throughout development^14^. Although we have not performed transcriptomic analyses in this study, the positive correlation between number of EdU+ cells, and growth (determined by increasing worms’ area) and gross development in HS-developed parasites was evident already within the first 48 hours in culture. These findings indicate that stem cells may play a central role in driving organised growth, tissue differentiation and ultimately the establishment of the sexual dimorphism in HS-developed parasites. Conversely, FBS-cultured parasites displayed no more than 20 EdU+ cells per worm in average, reaching a plateau in the number of proliferating cells from day 8 in culture, which probably underlie the arrested development of these worms. Overall, the FBS-cultured parasites did not progress beyond the lung stage. Consistent with these findings, confocal microscopy imaging revealed in HS-developed parasites clear somatic and reproductive anatomical structures, including male gynaecophoric canals and testes with sperm cells, as well as female ovary primordia, oviducts, ootypes and uteri. These developmental features illustrate the capacity of our culture system to capture key biological transitions previously only accessible from experimentally infected animal models. That said, while our system was highly efficient in producing sexually dimorphic worms, spontaneous pairing between male and female parasites was extremely rare, mainly in aged in vitro cultures (from 80 to 100 days in culture) indicating that other factors, e.g., cholesterol, may be missing^35^. In any case, we decided to test the pairing capacity of the dimorphic worms by conducting pairing experiments between *in vitro*-developed females and *ex vivo*-collected males, and vice versa. Strikingly, couples were observed within 24 hours after co-cultivating *in vivo* and *ex vivo* opposite sex parasites, albeit no eggs were produced. However, we observed evidence of pairing-trigger oocyte maturation in *in vitro*-developed females paired with *ex vivo*-collected male worms. Our research group has recently been involved in the discovery and functional characterisation of a transcription factor of the retinoic acid receptor family, SmRAR, and related genes^69^. SmRAR and associated genes may play a key role in oocyte differentiation triggered by pairing. Our pairing experiments and confocal imaging analyses suggested that *in vitro* developed female oocytes started to differentiate after pairing with an *in vivo* developed male worm, probably mediated by the activation of the SmRAR pathway^69^. Moreover, the involvement of a male-derived nonribosomal peptide pheromone, i.e., β-alanyl-tryptamine or ‘BATT’, in the pairing-driven female sexual maturation was demonstrated^6^. In addition, soluble factors produced by the worms with effect on the opposite sex cannot be ruled out^7^. It has been suggested that host-derived factors, such as the cytokine transforming growth factor β (TGF-β) may be critical for female reproductive development and embryogenesis^70^. More recently, other host-derived factors that include cholesterol and ascorbic acid have been shown to be critical for maintaining *ex vivo* fully mature females collected from infected mice^35^. Considering these elements in future experiments will help overcome the limitations encountered in this study, including the low rate of spontaneous pairing between *in vitro*–developed male and female worms and the requirement for extended culture periods (>70 days). In addition, further research is needed to assess the role of host- and parasite-derived cues in schistosome development^71^.

In summary, we have demonstrated that the presence of HS is essential to support fully *in vitro* development of *S. mansoni* parasites from mechanically transformed cercariae to sexually dimorphic adults. Differences in the numbers of proliferating stem cells between worms developed in HS versus FBS were observed as early as 48 hours in culture. These findings may raise some concerns about studies that rely on *in vitro* culture protocols using FBS, including those focused on parasite developmental biology, interaction with the host, and drug screening. Human serum contains essential host-specific molecules that are absent or insufficient in FBS, and future efforts should aim to identify these factors. Optimising schistosome long-term culture systems will: (1) deepen our understanding of fundamental aspects of schistosome biology, development and host-parasite molecular crosstalk; (2) positively impact the 3Rs principles for animal research; and (3) reveal tentative targets for novel control strategies.

## Material and Methods

### Ethics statement

The complete life cycle of *Schistosoma mansoni* (NMRI strain) was maintained at the Wellcome Sanger Institute (WSI) and Aberystwyth University (AU), by infecting *Biomphalaria glabrata* snails and outbred mice (TO strain). All the animal regulated procedures were conducted under Home Office Project Licences P77E8A062 (WSI) and PP2955700 (AU), following the ARRIVE guidelines (https://arriveguidelines.org) and in accordance with guidelines and regulations stated by the UK Animals (Scientific Procedures) Act 1986 Amendment Regulations 2012. All the protocols were presented and approved by the Animal Welfare and Ethical Review Bodies (AWERB) of the WSI and AU. The AWERB is constituted as required by the UK Animals (Scientific Procedures) Act 1986 Amendment Regulations 2012. Human blood products, including serum and red blood cells were obtained from NHS Blood and Transplant (NHSBT), Non-Clinical Issue (NCI) services for research purposes. NHSBT-NCI provides donated material surplus to clinical requirements or unsuitable for therapeutic use that has been appropriately consented. The supply chain complies with all statutory and regulatory obligations including (but not limited to) the Human Tissue Act (2004) and associated Codes of Practice. The NHSBT-NCI products were acquired under the Study ‘*In vitro* development of the human parasite Schistosoma’, Integrated Research Application System (IRAS) project ID 319400, protocol number AU/DLS/011, Research Ethics Committee (REC) reference number 23/SS/0017.

### Cercariae transformation and culture of schistosomula

*Schistosoma mansoni* (NMRI strain) schistosomula were obtained and cultured as previously described with minor modifications^19^. In brief, patent mix strain of *Biomphalaria glabrata* snails^72^ experimentally infected *en masse* with ∼20 miracidia per snail, were thoroughly rinsed, transferred to Lepple water (∼100ml) and exposed to light for 2 h at 26 °C to induce cercarial shedding. Cercariae were collected, passed through a 100μm filter into 50 ml tubes to remove snail debris and faeces, placed 1 h on ice, and concentrated by centrifugation (300 g for 3 min at 4°C with half break deceleration). Cercariae were washed 3 times in 1X PBS supplemented with 200 U/ml penicillin, 200 μg/ml streptomycin, 500 ng/ml amphotericin B (Merck). During the washes, the cercarial pellets were successively combined and finally resuspended in ‘Schistosomula wash medium’ (Dulbecco’s modified Eagle medium - DMEM, supplemented with 10 mM Hepes, 200 U/ml penicillin, 200 g/ml streptomycin, 500 ng/ml amphotericin B). Cercariae were vortexed full speed for 30 sec and placed on ice for 1 min. Thereafter, the cercarial tails were sheared off by ∼10 back and forth passes through a 19-gauge (19-G) needle, a ∼3 μl aliquot of parasites inspected under microscope to confirm that >90% of cercariae had their tail removed, and the cercarial heads were separated from the tails by a Percoll (Merck) gradient (1.5:1 percoll:DMEM) and centrifugation (300g for 15 min at 4 °C, half break deceleration). The pellet containing purified cercarial heads, was collected and washed 3 times by centrifugation (300g for 3 min at 4°C) in ‘Schistosomula wash medium’. Newly transformed schistosomula were counted in 12 5μl-aliquots under the microscope. To reduce parasite mortality due to a high density of worms, ∼8,000 schistosomula were transferred to each well of a 6-well tissue culture plate (Fisher Scientific, Loughborough, UK) containing 5 ml of modified Basch’s medium18 supplemented with additives and FBS or HS heat-inactivated FBS or HS serum, i.e., ‘Long-term culture medium’ (LTC) as indicated in Table 1. The number of parasites cultured per well (∼8,000 schistosomula) was determined empirically, as no formal titration experiments were performed. At higher densities (>10,000 per well), more frequent media changes were required, and parasite development appeared to be impaired. Parasites were cultured in a tissue culture incubator at 37°C under 5% CO2 in air. LTC medium was replaced twice a week and washed human red blood cells (hRBCs) added to a final concentration of 0.02% v/v at day 13 after transformation. Washed hRBCs were replaced every two weeks, or sooner if their numbers decreased due to consumption.

**Table 1.**
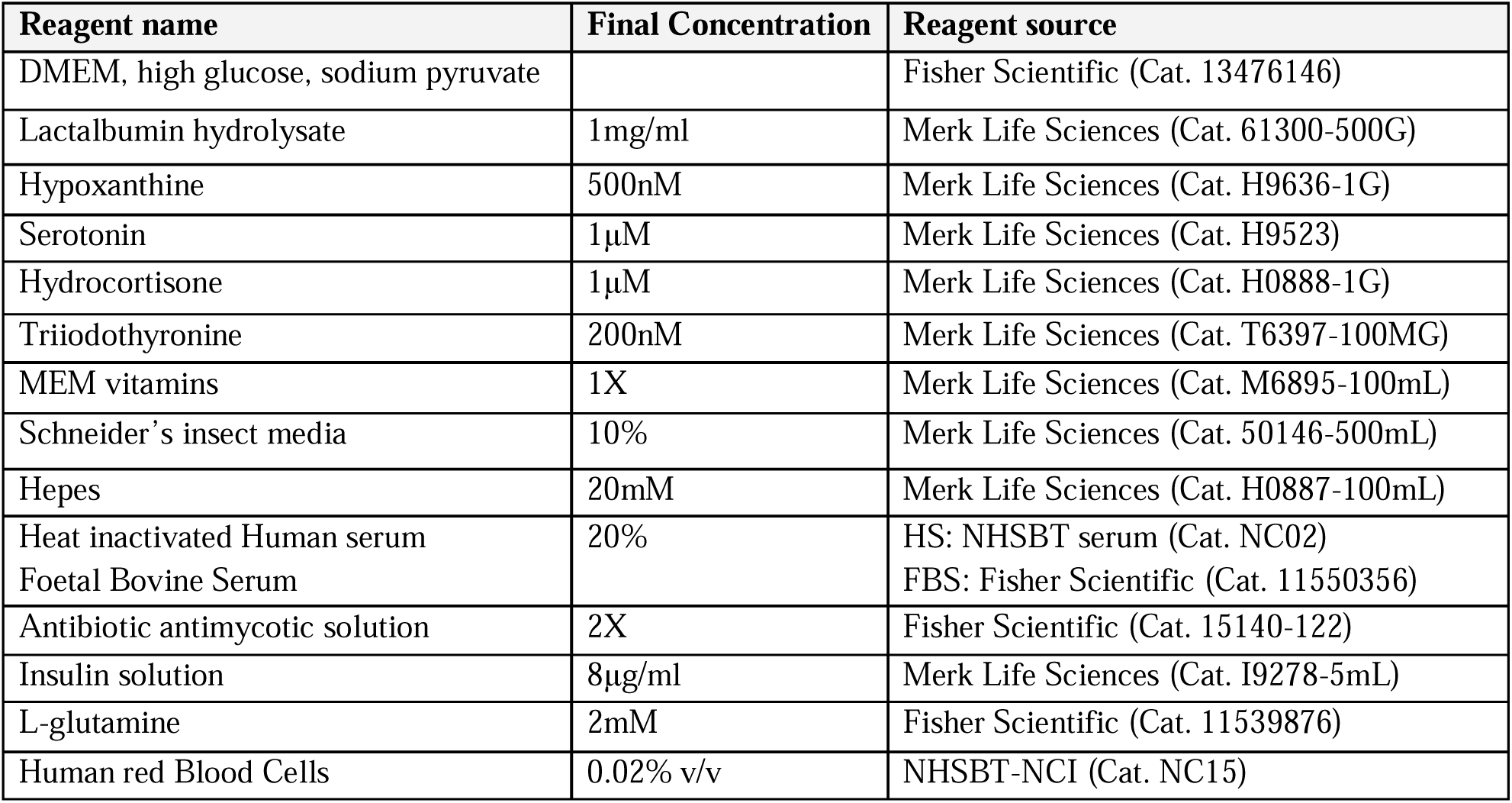
Long term culture medium (LTC medium) composition.

All human blood products, obtained from NHSBT-NCI, were handled with universal precautions within a biosafety cabinet class II and suitable PPE. The HS was heat-inactivated at 56°C for 30 min in an incubator, aliquoted and stored at −20°C until use. The hRBC were washed in ‘Schistosomula wash medium’ and stored at 4°C until use as previously described^19^. Briefly, the hRBCs were transferred from the pack to 50 ml tubes and washed by centrifugation at 500g for 5 min at 4°C. Half the supernatant was removed, tubes’ content combined, centrifuged as above and resuspended in ∼25 ml of ‘Schistosomula wash medium’. The washes were repeated 4 more times.

### Phenotype scoring of developing schistosomula

To analyse the *in vitro* development of schistosomula, images and videos were taken at regular intervals of at least once a week in >10 independent culture experiments, using a digital Euromex camera connected to an Olympus CK2 optical microscope. Up to 18 pictures per experiment with an average of ∼400 worms per picture were used for parasite staging from at least 5 independent experiments (**Supplementary Table S1**). The investigators who assigned categories were blinded to experimental conditions. Based on a well-defined staging system^12,33^, the parasites were classified into 6 developmental categories as described^39^ (**Supplementary Figure S1**); category 0: dead parasites showing granulation, rounded shape and degraded or broken tegument - the number of dead parasites may be underestimated due to the loss of some dead worms during the media change; category 1: newly transformed schistosomula, corresponding to the *in vivo* ‘skin stage’, these worms retain the cercarial head shape with no visible internal structures such as gut; category 2: lung stage schistosomula, elongated and slim worms, usually with a bulged end; category 3: early liver stage, bigger and wider worms compared to the previous stage, with hemozoin pigment (i.e., haemoglobin degradation product) and two ceca gut; category 4: late liver stage, in which the gut is further developed, the two intestinal ceca have fused behind the ventral sucker, and the parasites started to acquire an evident vermiform shape; category 5: sexually dimorphic stage, comprising large, vermiform male- and female- looking worms. The females are longer and thinner than males, the ceca fusion localises closer to the anterior end of the worm (i.e., ∼⅓ of the worm length from the anterior end), and both suckers are smaller compared to the male suckers. The male worms are characterised by a larger and wider head, larger suckers, the ceca fusion closer to the posterior end of the animal (i.e., ∼⅔ of the worm length from the anterior end), and a clearly visible gynecophoral groove^12,33^. To evaluate differences in mortality between HS- and FBS-cultured parasites, data from 5 experiments were combined and analysed using a Shapiro-Wilk normality test to test normality of the data and a non-parametric Wilcoxon rank sum exact test (**Supplementary Tables S1 and S6**).

To quantify positive or negative parasites for the presence of hemozoin within their intestines, i.e., black guts + (BG+) or BG- parasites, respectively, hRBCs were added to the culture at day 13, and one or two days later bright field microscopy images were taken. Counts of BG+ and BG- parasites were collected from 5 independent *in vitro* culture experiments (**Supplementary Table S4**). Statistical analysis was performed using a Wilcoxon rank-sum (Mann-Whitney) test, with P≤0.05 considered significant (**Supplementary Table S6**).

### Measurement of parasite area

Images of *in vitro* developing parasites were taken at weeks 1, 2, 3, 4, 6, 8 and 10 after cercariae transformation with an Euromex camera fitted to an Olympus CK2 optical microscope at 4x and 10x magnification. For these magnifications images of scale bars were captured for pixel to area values (µm²). Parasites from 5 independent long-term culture experiments were masked in COCO format from captured images using the CVAT (Computer Vision Annotation Tool) online server (https://www.cvat.ai/). Area metrics were then calculated for each parasite by leveraging the pycocotools package in a customised Python script. Raw and processed data is provided in **Supplementary Table S3**. Normal distribution was checked using Shapiro test and Mann Whitney tests performed with a significant value set up for P≤0.05 (**Supplementary Table S6**).

### PCR for sexing and confirmation of sexual dimorphism

To confirm the *in vitro* establishment of sexual dimorphism, parasites were individually collected from culture by day ∼80, for blind sex assignment by bright field microscopy and confirmation by sex-specific PCR^73^ (**Supplementary Figure S3B**). Briefly, pictures from individual *in vitro* developed parasites were taken and sex assigned based on morphological features. Thereafter, genomic DNA from the sex-assigned individual parasites was extracted by lysing the worms at 95°C for 1h in 50mM NaOH, 0.4mM disodium ethylenediaminetetraacetic acid (EDTA), followed by the addition of neutralising buffer (80 mM Tris–HCl in water). Samples were stored at 4°C until use. PCR reactions were performed in a final volume of 10μl consisting of 5μl Platinum 2X hot start PCR master mix (Invitrogen, California, USA), 0.5μl (10μM stock) of forward and reverse primers targeting W1 repeat^74^ and actin^75^ (**Supplementary Table S2**), 1μl of template DNA and 3μl of water. The PCR program comprised an initial denature step at 94°C for 2 min followed by 30 cycles of denature at 98°C for 5 secs, and annealing/elongation at 60°C for 15 secs. The PCR products were resolved on a 1% agarose gel using 1% TAE buffer and 1kb GeneRuler DNA ladder (Thermo Scientific, UK).

### Detection and quantification of proliferating cells

Parasites cultured for 15 days (D15) in media supplemented with either HS or FBS and hRBC added at D13 were harvested at days 2, 8 or 15 after cercariae transformation. Parasites cultured under the same conditions but with no hRBC were included as controls. The day before each indicated time point, a solution of 5-ethynyl-2’-deoxyuridine (EdU) was added to the culture medium at 10 μM final concentration. At each indicated time point, parasites from each group were collected and transferred to a 35 µm mesh basket (CEM Microwave Tech) in a well of a 24-well plate, washed in 1x PBS, fixed in 4% Paraformaldehyde (PFA) in 1x PBS for 30 min, washed in 1x PBS and stored at 4°C until use. The worms were incubated in PBST (1x PBS + 0.3% Triton x-100) containing 2μl of Proteinase K (20 mg/ml) for 5 min at room temperature, washed three times in PBST, fixed in 4% PFA in 1x PBS for 10 min at room temperature and washed 5 times in PBST. EdU-positive cells were revealed by incubating the parasites for 30 min in the dark in a solution containing Azide fluor 488 (Sigma-Aldrich), diluted in a solution of 1mM CuSo4, 0.025mM Azide fluor 488, 95mM L-Ascorbic Acid in 1x PBS. The worms were washed 5 times in PBST and incubated in mounting media containing DAPI (Fluoromount-G™ Mounting Medium, with DAPI, Invitrogen) before mounting for confocal microscopy (Leica SP8 super resolution laser confocal microscope). EdU+ cells per parasite were counted for an average of 100 parasites across three independent experiments (**Supplementary Table S4**). Worms were grouped based on the number of cells per individual, but all those showing Ill60 EdU+ cells were counted in the same group named ‘60 EdU+ cells’. Therefore, the data were considered ordinal and the statistical analysis performed by Kruskal-Wallis test with Dunn multiple comparison post-hoc test, with P≤0.05 considered significant (**Supplementary Table S6**).

### Pairing experiments

*Schistosoma mansoni* adult male and female worms were collected from experimentally infected mice at 47 days post infection^19^. Briefly, mice were euthanised by intraperitoneal injection of 200 μl of 200 mg/ml pentobarbital supplemented with 100 U/ml heparin, and worms collected by portal perfusion (the hepatic portal vein was sectioned followed by intracardiac perfusion with phenol-red-free DMEM, containing 10 U/mL heparin) and washed in DMEM. In 24-well plates, *in vitro* developed female worms were cultured in the presence of *in vivo* developed male worms and vice versa at a ratio of 1 female every 2 males in the well containing culture medium (**Table 1**) supplemented with 20% HS. The parasites were cultured for several days in an incubator at 37°C, 5 % CO2 and checked daily under microscope for the presence of pairing.

### Parasite staining and confocal imaging

Live parasites collected from culture at indicated time points were stained with 1:1000 dilutions of CellMaskTM Green Actin Tracking Stain (Invitrogen) and CellMaskTM Deep Red Plasma Membrane Stain (Invitrogen) to label polymerized/filamentous actin (F-actin) and cell membranes, respectively. Nuclei were stained by adding two drops per millilitre of NucBlueTM Live ReadyProbesTM Reagent (Invitrogen). After an overnight incubation at 37°C, worms were collected in 15 ml tubes, washed by gravity 3 times in 1x PBS, fixed in 4% PFA (in 1x PBS) overnight at 4°C, washed 3 times in 1x PBS and stored in 200 μl of Fluoromount-G, with DAPI (Invitrogen) before mounting on slides for confocal microscopy. All the images were acquired using a Leica SP8 super resolution laser confocal microscope.

## Supporting information

Supplementary TablesR1

## Acknowledgements

The authors thank Julie Hirst for technical assistance with the *Schistosoma mansoni* life cycle maintenance at Aberystwyth University, and Anais Bordes for technical support with the *Schistosoma mansoni* life cycle maintenance at University of Oxford. The authors also gratefully acknowledge Jennifer Holter from the Biology Dept Imaging Suite, University of Oxford for their support & assistance in this work (Funder: EPA Cephalosporin Fund and Department of Biology, Project Number CBR00830/REF CF 401). The authors acknowledge NHS Blood and Transplant (NHSBT), Non-Clinical Issue (NCI) for providing human blood products (detailed information in Methods). The study was partially funded by the Wellcome Trust (grants 098051 and 206194), and the European Union [Project 101080784 – WORMVACS2.0]. GR is supported by UKRI Future Leaders Fellowships [MR/W013568/2].

## Author contributions

Remi Pichon, Conceptualization, Methodology, Investigation, Data curation, Formal analysis, Visualization, Writing – original draft, Writing – review and editing; Magda E Lotkowska, Conceptualization, Methodology, Investigation, Data curation, Formal analysis, Visualization; Jude L. D. Bulathsinghalage, Methodology, Investigation, Data curation; Madeleine McMath, Methodology, Investigation, Data curation, Formal analysis, Visualization; Mary Evans, Methodology, Investigation; Benjamin J. Hulme, Methodology, Investigation; Kirsty Ambridge, Methodology, Investigation, Visualization; Geetha Sankaranarayanan, Methodology, Investigation; Simon Kershenbaum, Formal analysis, Writing – original draft, Writing – review and editing; Sarah D. Davey, Methodology, Investigation, Data curation, Formal analysis, Visualization; Josephine E. Forde-Thomas, Methodology, Investigation, Visualization; Karl F. Hoffmann, Conceptualization, Supervision, Funding acquisition; Matthew Berriman, Supervision, Funding acquisition, Project administration; Gabriel Rinaldi, Conceptualization, Methodology, Investigation, Data curation, Formal analysis, Visualization, Supervision, Funding acquisition, Project administration, Writing – original draft, Writing – review and editing.

## Supplementary information

### Supplementary Tables

**Supplementary Table S1.** Raw counts of parasites within each developmental stage category. Each row corresponds to a picture of parasites in culture medium containing FBS or HS. Each column corresponds to the raw parasite counts at indicated stage development (categories 0 to 5), time in culture (Time in days - D), and experimental condition.

**Supplementary Table S2.** Primers used in PCR for sexing parasites

**Supplementary Table S3.** Area raw measurements of developing worms

**Supplementary Table S4.** Percentage of parasites with either black positive (hemozoin) or black negative (no hemozoin) intestine. BG: Black gut.

**Supplementary Table S5.** Raw counting of EdU positive cells per parasite for indicated experimental group, replicate and experiment in long format. The worms were classified by group (column C) and replicate (column D), using the following code: E (‘early’), M (‘medium’) and L (‘late’), corresponding to days 2, 8 and 15, respectively. R and W correspond to conditions with (R) or without (W) human red blood cells, and HS and FBS to culture medium employed.

**Supplementary Table S6.** Summary of all statistical tests employed in this study. 1. Statistical tests of parasite mortality and the raw data table used for this test. 2. Statistical tests for worm size comparisons (correspond to Figure 2). 3. Statistical tests for worm black gut comparisons (correspond to Figure 3). BG: Black gut. 4. Statistical tests for EdU positive cells comparisons (correspond to Figure 4). Replicate code: E, M and L correspond to day 2, 8 and 15 respectively; R and W correspond to the presence (R) or absence (W) of RBCs added 13 days after transformation.

### Supplementary Figures

**Supplementary Figure S1.** Developmental staging scoring. Summary of morphological attributes for the five different categories of development of worms cultivated *in vitro* (HS), as well as representative bright field images of each of these morphological categories. Arrows in categories 2 and 4 indicate elongated lung schistosomula, and the fusion of the two ceca posterior to the ventral sucker in the late liver schistosomula, respectively. Scale bar: 100 µm.

**Supplementary Figure S2.** Representative FBS-cultured parasites by w eek 10 in culture; most of the worms are dead (red arrows) and the few still alive are lung schistosomula (green arrows). Scale bar: 100 µm.

**Supplementary Figure S3. A.** Representative pictures of *in vitro* developed parasites by day ∼80 in culture, male or female worms as indicated. Scale bar: 500 µm **B.** Representative results of the PCR-based sex genotyping using primers to amplify a fragment of the control actin gene in both male and female (upper band), and primers to amplify a W-specific region only in female worms (lower band). A non-template control, and male and female positive controls were included as indicated. Assigned sexes for each of the groups of clonal cercariae emitted by individual snails infected with single miracidia.

**Supplementary Figure S4.** Representative confocal high magnification images of individual parasites displaying Edu+ cells at each indicated time point and culture condition. Edu+ cells and nuclei were labelled with Alexa fluor 488 (green) and DAPI (white/grey), respectively. Scale bars: 50 µm.

**Supplementary Figure S5.** Representative bright-light pictures parasites cultured in HS supplemented medium for more than 100 days, indicating male and female worms. A-F: parasites cultured for 103 days. G, H. Parasites cultured for 145 days. Scale bar: 100 µm.

### Supplementary Videos

**Supplementary Video S1.** Representative Z-stack of EdU+ cells after 2 days of *in vitro* culture with human serum (A) or foetal bovine serum (B). EdU+ cells and nuclei were labelled with Alexa fluor 488 (green) and DAPI (grey), respectively. Scale bars: 50 µm.

**Supplementary Video S2.** Representative Z-stack of an *in vitro* developed male worm. CellMask Green Actin Tracking Stain (green), and DAPI-stained nuclei (cyan). Scale bar: 25 µm.

**Supplementary Video S3.** Representative Z-stack of an *in vitro* developed female worm. CellMask Green Actin Tracking Stain (green), and DAPI-stained nuclei (cyan). Scale bar: 10 µm.

**Supplementary Video S4.** Representative video of *in vitro* developed females and males *in copula* at ∼80 days in culture. Scale bar: 100 µm

**Supplementary Video S5.** Representative videos of *in vivo* developed male and *in vitro* developed female *in copula* (A), and *in vivo* developed female and in *vitro* developed male *in copula* (B) Scale bar: 100 µm

**Supplementary Video S6.** Z-stack of a schistosome pair (*in vivo-*developed male and *in vitro*-developed female) *in copula* (A), and magnification of the ovarian area (B), highlighting maturing oocytes. CellMask Green Actin Tracking Stain: green, DAPI: grey, CellMask Deep Red Plasma Membrane Stain purple. Scale bars: 25 µm (top panel) and 50 µm (bottom panel)

