## Supplementary Figures for "*In vitro* sexual dimorphism establishment in schistosomes"

Figure S1

| | | |
|---|---|---|
| **Category 0:** Dead schistosomula | - Usually 'roundish'<br>- Severe granulation<br>- "Dark granules in the body"<br>- Tegument NOT smooth, disrupted | 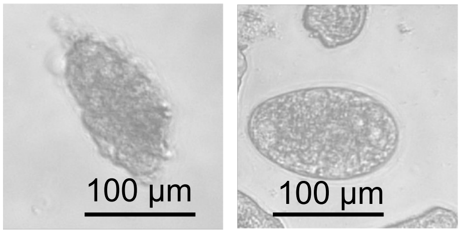 |
| **Category 1:** Early schistosomula | - Cercarial head shape<br>- Short, roundish, clear ('transparent')<br>- Gut not developed (no black pigment) | 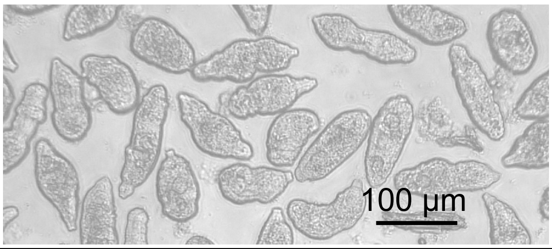 |
| **Category 2:** Lung schistosomula | - Elongated and thin<br>- No gut development (no black pigment)<br>- One end wider, roundish | 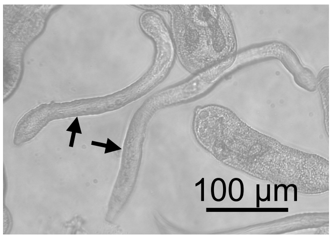 |
| **Category 3:** Early Liver schistosomula | - Bigger, wider<br>- Suckers more developed<br>- Gut started to develop (black pigment)<br>- Gut NOT YET fused behind the ventral sucker | 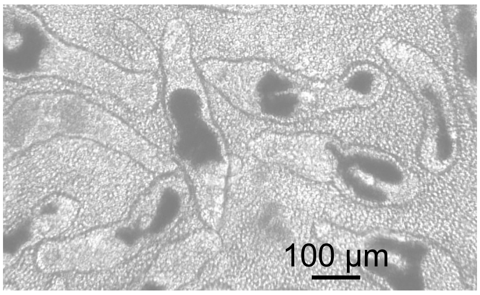 |
| **Category 4:** Late Liver schistosomula | - Bigger, wider<br>- Suckers more developed<br>- Gut FUSED behind the ventral sucker | 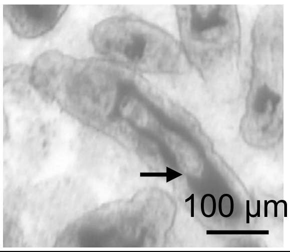 |
| **Category 5:** Dimorphic schistosomula | - Bigger, developed suckers<br>- Females longer, thinner, smaller head and suckers, 1/3 double-ceca – 2/3 single cecum<br>- Males wider but sometimes shorter than females, wider head and bigger suckers, 2/3 double-ceca – 1/3 single cecum | 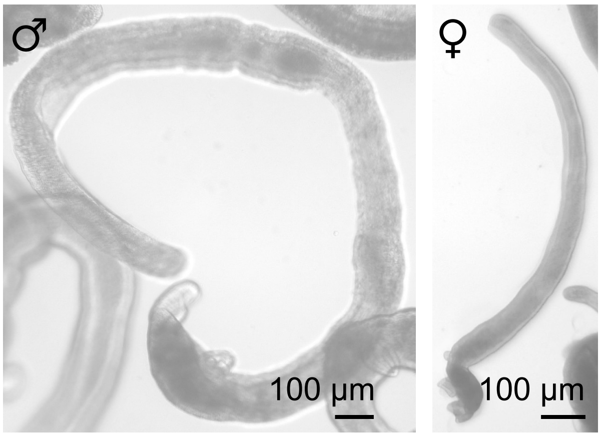 |

Figure S2

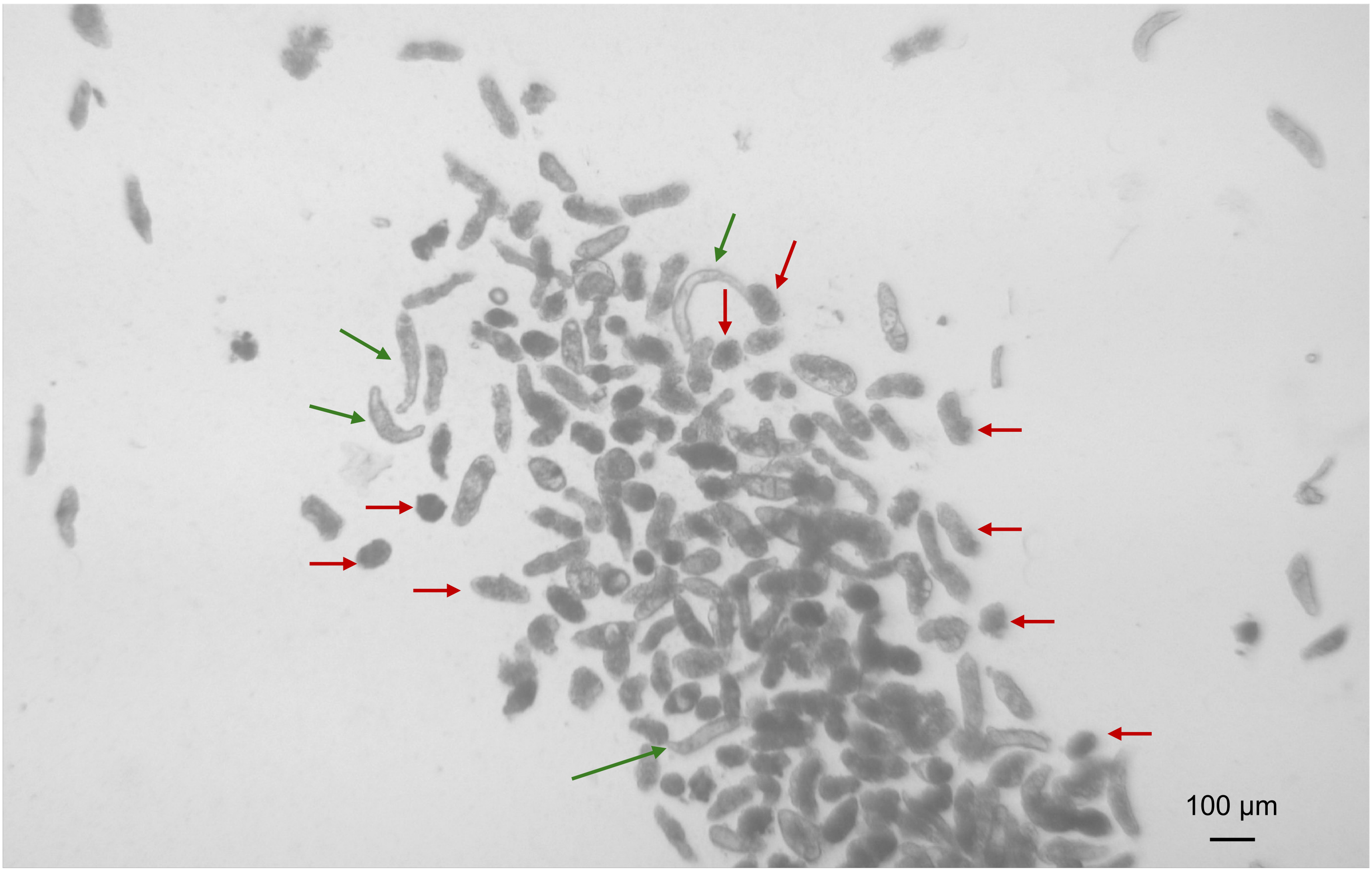

Figure S3

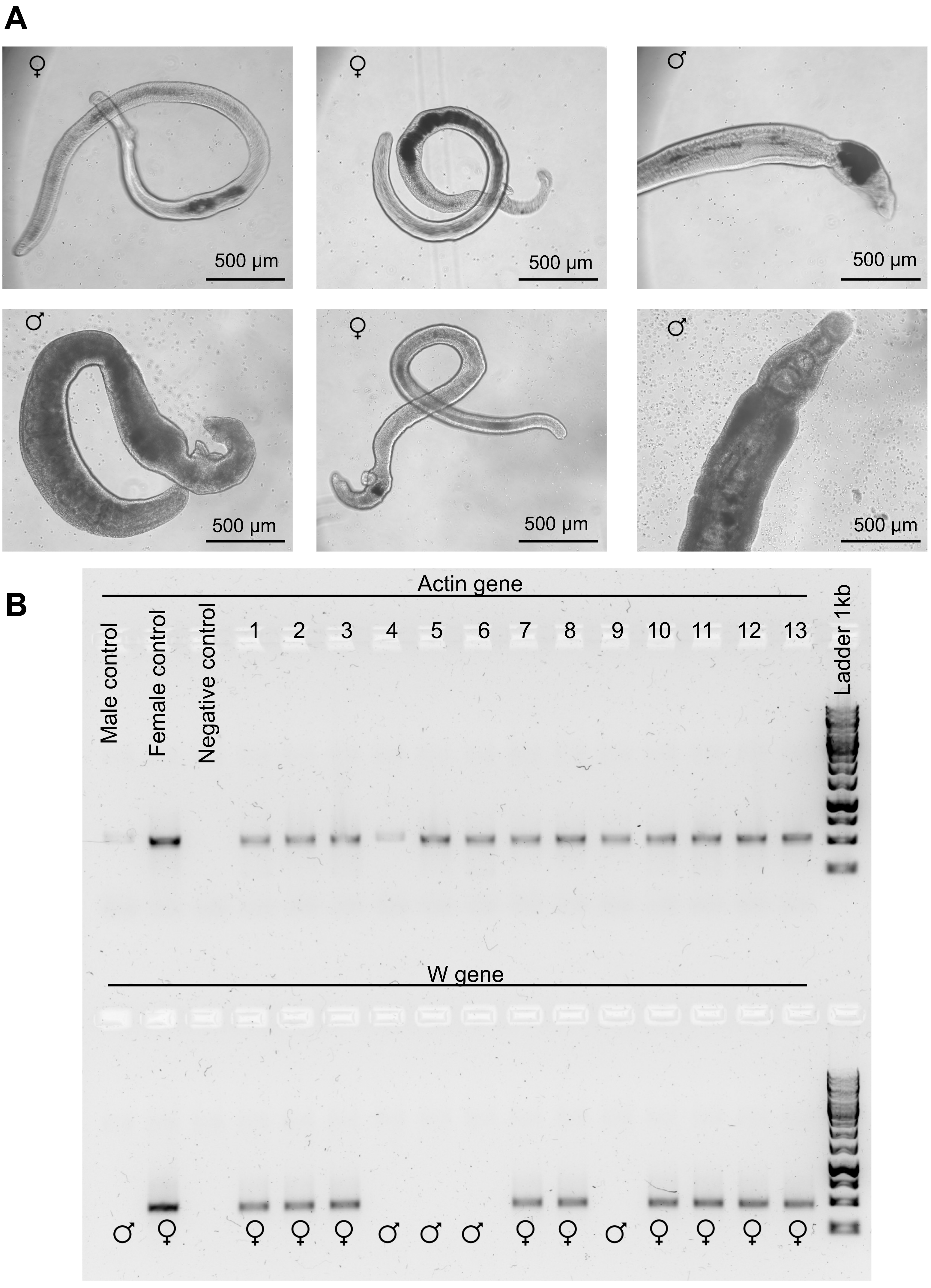
A
♀
500 µm
♀
500 µm
♂
500 µm
♂
500 µm
♀
500 µm
♂
500 µm
B
Actin gene
Male control
Female control
Negative control
1
2
3
4
5
6
7
8
9
10
11
12
13
Ladder 1kb
W gene
♂
♀
♀
♀
♀
♂
♂
♂
♀
♀
♂
♀
♀
♀
♀

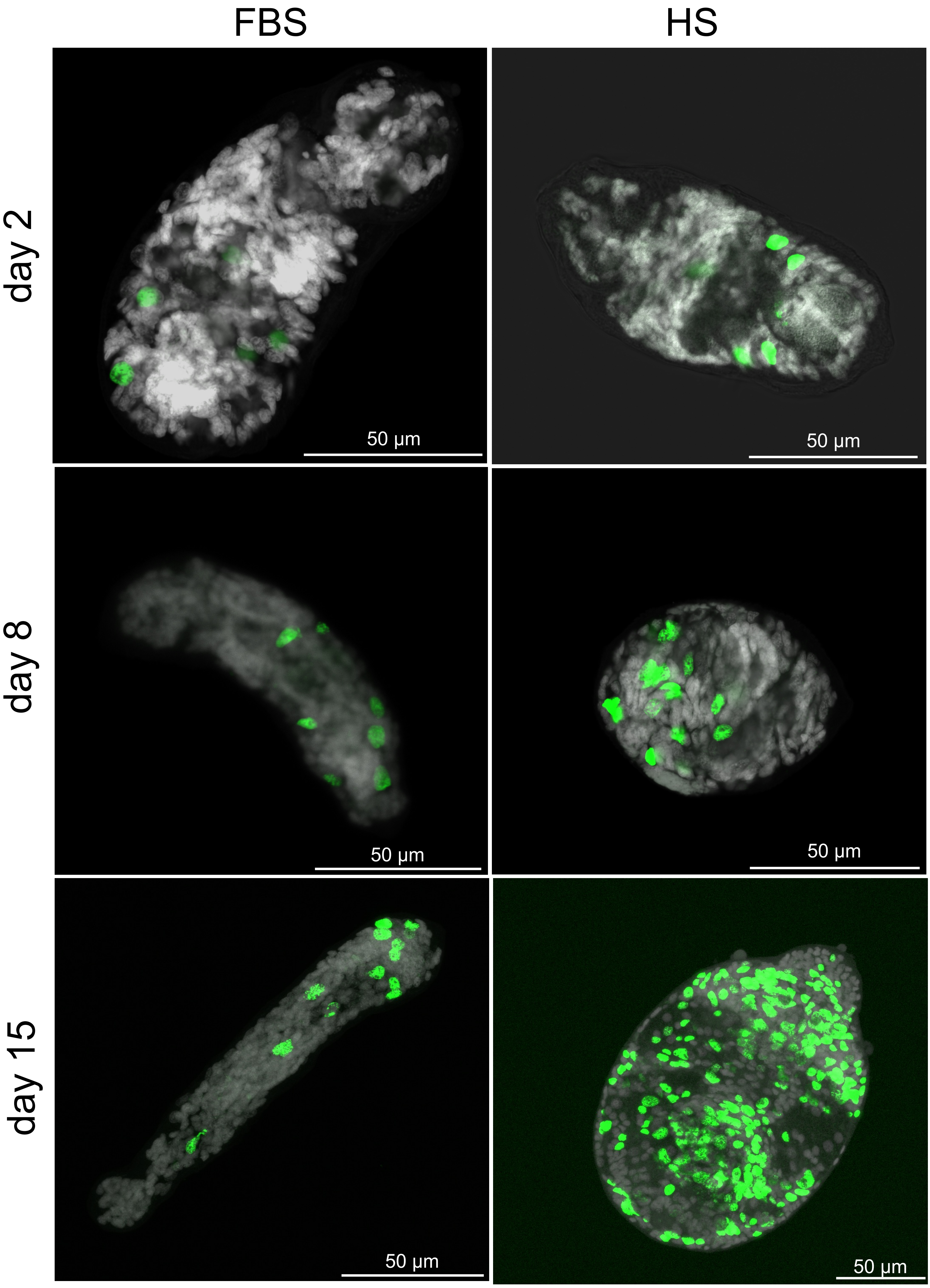
FBS
HS
day 2
day 8
day 15
50 µm
50 µm
50 µm
50 µm
50 µm
50 µm

Figure S5

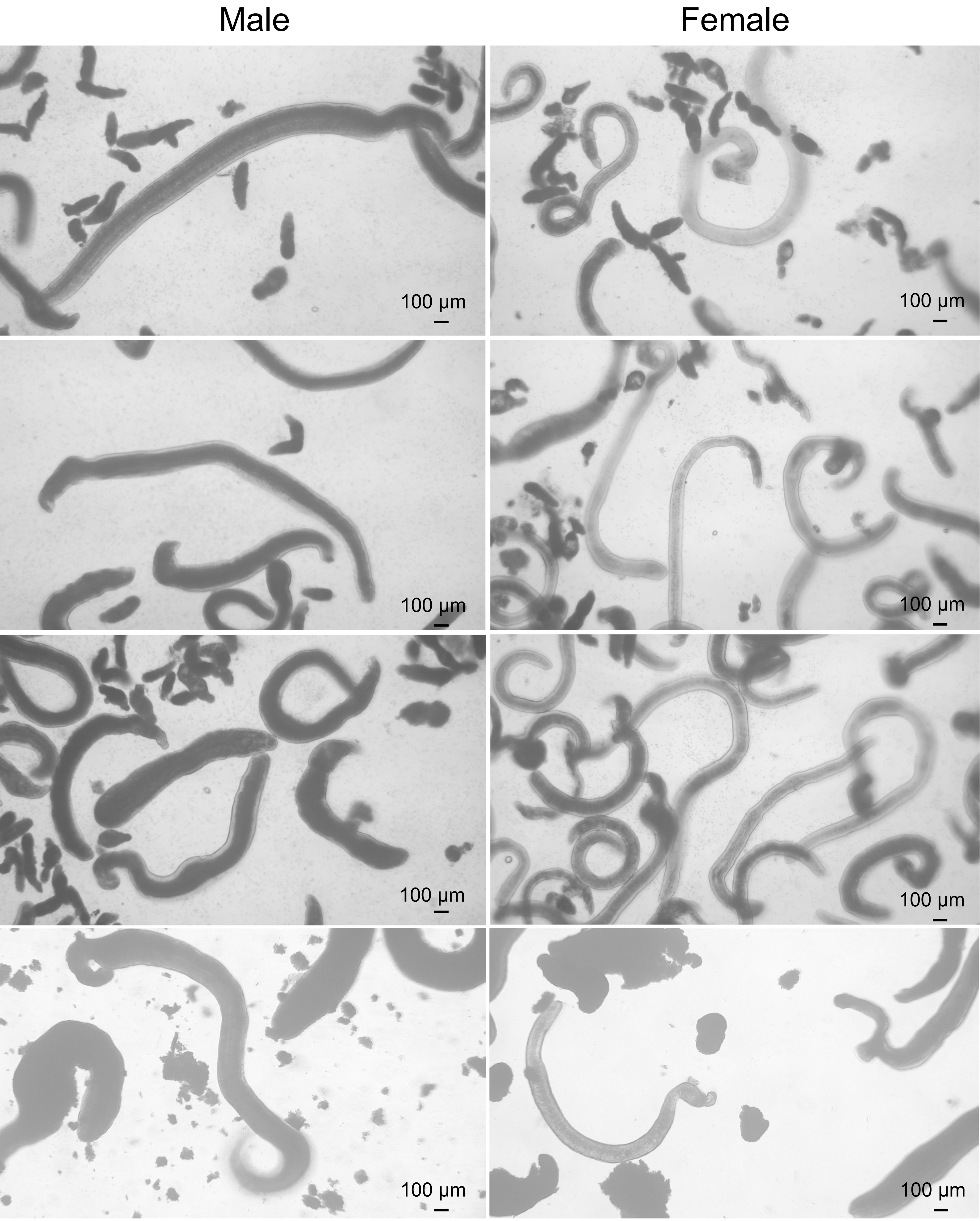
Male
Female
100 µm
100 µm
100 µm
100 µm
100 µm
100 µm
100 µm
100 µm
