## Supplementary Videos for "*In vitro* sexual dimorphism establishment in schistosomes"

### 1. Department of Biology, University of Oxford, Oxford, UK

### 2. Department of Life Sciences, Aberystwyth University, Aberystwyth, UK

### 3. Wellcome Sanger Institute, Hinxton, UK

### 4. School of Infection and Immunity, College of Medical, Veterinary & Life Sciences, University of Glasgow, Glasgow, UK

*co-first authors, **co-corresponding authors

Correspondence:

Matt Berriman:

Gabriel Rinaldi:

**Supplementary Videos**

**Supplementary Video S1.** Representative Z-stack of EdU+ cells after 2 days of *in vitro* culture with human serum (A) or foetal bovine serum (B). EdU+ cells and nuclei were labelled with Alexa fluor 488 (green) and DAPI (grey), respectively. Scale bars: 50 µm.

<https://www.dropbox.com/scl/fi/af487tikkdvpjy5kxba9r/Pichon-Lotkowska_etal_VideoS1_BioRxiv.pptx?rlkey=capqp6yv5gvibglkmhkm8l0n1&st=c98cv9np&dl=0>

**Supplementary Video S2.** Representative Z-stack of an *in vitro* developed male worm. CellMask Green Actin Tracking Stain (green), and DAPI-stained nuclei (cyan). Scale bar: 25 µm.

<https://www.dropbox.com/scl/fi/fi5rg6m1hm530si164vrt/Pichon-Lotkowska_etal_VideoS2_BioRxiv.pptx?rlkey=ou8a8jq7a1vyihrqdnagnyj9j&st=zq60jrd1&dl=0>

**Supplementary Video S3.** Representative Z-stack of an *in vitro* developed female worm. CellMask Green Actin Tracking Stain (green), and DAPI-stained nuclei (cyan). Scale bar: 10 µm.

<https://www.dropbox.com/scl/fi/axihef0arhffj18s53rol/Pichon-Lotkowska_etal_VideoS3_BioRxiv.pptx?rlkey=60d2wp5rutz7oflwofazrewnv&st=j5xk517j&dl=0>

**Supplementary Video S4.** Representative video of *in vitro* developed females and males *in copula* at ~80 days in culture. Scale bar: 100 µm

<https://www.dropbox.com/scl/fi/dyk1fyzrhxvupq2lbriax/Pichon-Lotkowska_etal_VideoS4_BioRxiv.pptx?rlkey=4lk6zgpf9uea253uivrdcsq1i&st=cpxkb9ue&dl=0>

**Supplementary Video S5.** Representative videos of *in vivo* developed male and *in vitro* developed female *in copula* (A), and *in vivo* developed female and in *vitro* developed male *in copula* (B) Scale bar: 100 µm

<https://www.dropbox.com/scl/fi/952ucod7so6fj38x5g4d7/Pichon-Lotkowska_etal_VideoS5_BioRxiv.pptx?rlkey=13ost4m9mffznnzu6lkd66fup&st=rcy99fdp&dl=0>

**Supplementary Video S6.** Z-stack of a schistosome pair (*in vivo-*developed male and *in vitro*-developed female) *in copula* (A), and magnification of the ovarian area (B), highlighting maturing oocytes. CellMask Green Actin Tracking Stain: green, DAPI: grey, CellMask Deep Red Plasma Membrane Stain purple. Scale bars: 25 µm (A) and 50 µm (B)

<https://www.dropbox.com/scl/fi/yfamy9idmoxldzw9yn4t8/Pichon-Lotkowska_etal_VideoS6_BioRxiv.pptx?rlkey=ofe7s1hmltq378kgt8nigxnbk&st=404lvej6&dl=0>
